# Distinguish characters of luminal and glandular epithelium from mouse uterus using a novel enzyme-based separation method

**DOI:** 10.1101/2022.08.27.505528

**Authors:** Qi-Xin Xu, Wang-Qing Zhang, Lei Lu, Ke-Zhi Wang, Ren-Wei Su

## Abstract

The uterine luminal epithelium, glandular epithelium, and stromal cells are vital for the establishment of pregnancy. Previously studies have shown various methods to isolate mouse uterine epithelium and stromal cells, including Laser Capture Microdissection (LCM), enzyme digestion, and Immunomagnetic beads. Despite the importance of the endometrial epithelium as the site of implantation and nutritional support for the conceptus, there is no isolated method to separate the luminal epithelium and glandular epithelium. Here, we establish a novel enzyme-based way to separate two types of epithelium and keep their viability. In this article, we analyzed their purity by mRNA level, immunostaining, and transcriptome analysis. Our isolation method revealed several unstudied luminal and glandular epithelial markers in transcriptome analysis. We further demonstrated the viability of the isolated epithelium by 2D and 3D cultures. The results showed that we successfully separated the endometrial luminal epithelium and glandular epithelium. We also provided an experimental model for the following study of the physiological function of the different parts of the uterus and related diseases.

## Introduction

The uterus is an organ critical for mammalian reproduction, providing a proper micro-environment and nutrition supply to developing embryos and fetuses to achieve pregnancy success [1]. The uterine endometrium contains many types of cells, including endothelial cells, immune cells, stromal cells, and epithelial cells [2]. The diverse characters and functions of these cell types and their temporal and spatial interaction in the endometrium are fundamental to the progressive processes of the physiological functions of the uterus [3]. In addition, the epithelial cells can be further divided into two distinguished types: luminal epithelial cells (LE) and glandular epithelial cells (GE). LE and GE distinguish location, morphology, physiological function, hormone response, and differentiation potentials [4].

The LE cells are tall columnar, cover the uterine lumen’s surface, and serve as a barrier to separate the sperm, embryo and other contents from the parts of the endometrium. LE structural changes are prerequisites for establishing transient uterine receptivity for embryo implantation [5]. During a short period named the window of implantation (WOI), in response to ovarian hormones estrogen and progesterone, the LE cells change their morphology to short columnar or cuboidal, loss their polarity, and become receptive to the implantation of the blastocyst [6]. In contrast, the GE cells are cuboidal. The GE cells formed endometrial glands extend from the luminal epithelium into the surrounding stroma. They produce nutrition supply to embryos and key factors that control implantation, such as the secretion of Leukemia Inhibitory Factor (LIF) [7]. Uterine glands are derived from the luminal epithelium [8]. In mice, the development of endometrial glands begins postnatally, involving the differentiation and invagination of epithelial cells from luminal epithelium into the sub-surrounding stroma, through the processes including the budding of epithelial cells, extensive tubular coiling, and branching morphogenesis throughout the uterine stroma towards the myometrium [9]. On the other hand, in adults, evidence suggests that some cells in the gland may act as stem/progenitor cells that can regenerate both LE and GE during estrus and pregnancy cycles in humans and mice [10]. Therefore, collecting and studying LE and GE cells separately is key to understanding the different characteristics and functions of LE and GE cells.

In 1980 Glasser SR et al. developed an enzymatic digestion method to separate the epithelium from the longitudinally slit uterus of immature female rats [11, 12]. This method was then expanded to the mouse uterus by Kover K et al., and the remaining tissue after epithelium isolation was further digested to isolate stromal cells [13]. This method and its modified versions have been used in many studies for decades [14-16]. However, most of the epithelial cells isolated by this method from the mouse uterus are LE cells as they are TROP2 positive [17].

On the other hand, in 1986, Cunha GR and co-workers reported a simple, efficient, mechanical separation method that is able to isolate the intact epithelium from neonatal and older mouse uterus using a Pasteur pipette [18]. This method can obtain the pure intact luminal epithelium for the neonatal uterus as the glands are not genesis yet. For the glands developed stage, like 20 days of age, the epithelium isolated by this method comprises both LE and GE [18]. We have previously isolated the LE of E3.5 mice using a modified approach of this method, resulting in LE collection with limited GE contamination [19]. Nonetheless, to our best knowledge, no researchers focus on isolating GE fraction from the remaining tissue after LE isolation. In contrast to the mice case, the enzymatic method separating endometrial epithelial and stromal cells usually obtains more GE fraction than LE due to the lack of the LE layer during the sample collection process in humans [20, 21]. Nonetheless, none of the above methods is able to separate the LE and GE from the mouse uterus.

Filant J et al. applied Laser Capture microdissection (LCM) to isolate the GE and LE fraction from the mouse uterus and compared the transcriptomes of GE and LE cells on postnatal day 10, E2.5, and E3.5 using microarray [22]. However, the cells isolated by LCM can only be used for detecting the expression of molecules such as mRNA and protein, even by high-throughput technologies. The need for cell culture in further investigations regarding the difference between LE and GE required a method to separate these fractions from the uterus and keep them alive for in vitro culture.

In this study, inspired by the isolation method of human endometrial cell fractions, we developed the current method base on modified Cunha’s method: firstly, ensure the separation of intact LE without contamination of GE or S cells via modified Cunha’s method, and secondly, separate the GE fraction from the remaining GE-S combination by collagenase. We have successfully separated the LE, GE, and S fractions and identified their purities by expression of specific markers. The viability of cells was verified via 2D and 3D cultures. We further analyzed the difference in transcriptome between the LE and GE fractions and compared it to the LCM data.

## Results

### 1. Research Strategy

Compared to the coiled, branched glandular epithelium (GE) structure, the luminal epithelium (LE) cells form a monolayer on the internal surface of the uterine endometrium, which would be easier to be isolated intactly. Therefore, we designed our strategy as shown in Fig. 1: First, we removed the intact uterine luminal epithelium without contact with any uterine gland; then, we chopped the remaining fractions of the uterus, including the glands, stroma, and myometrium, into small pieces, and digested by collagenase to remove surrounding stromal cells and collect the pure glands.

**Fig 1.**
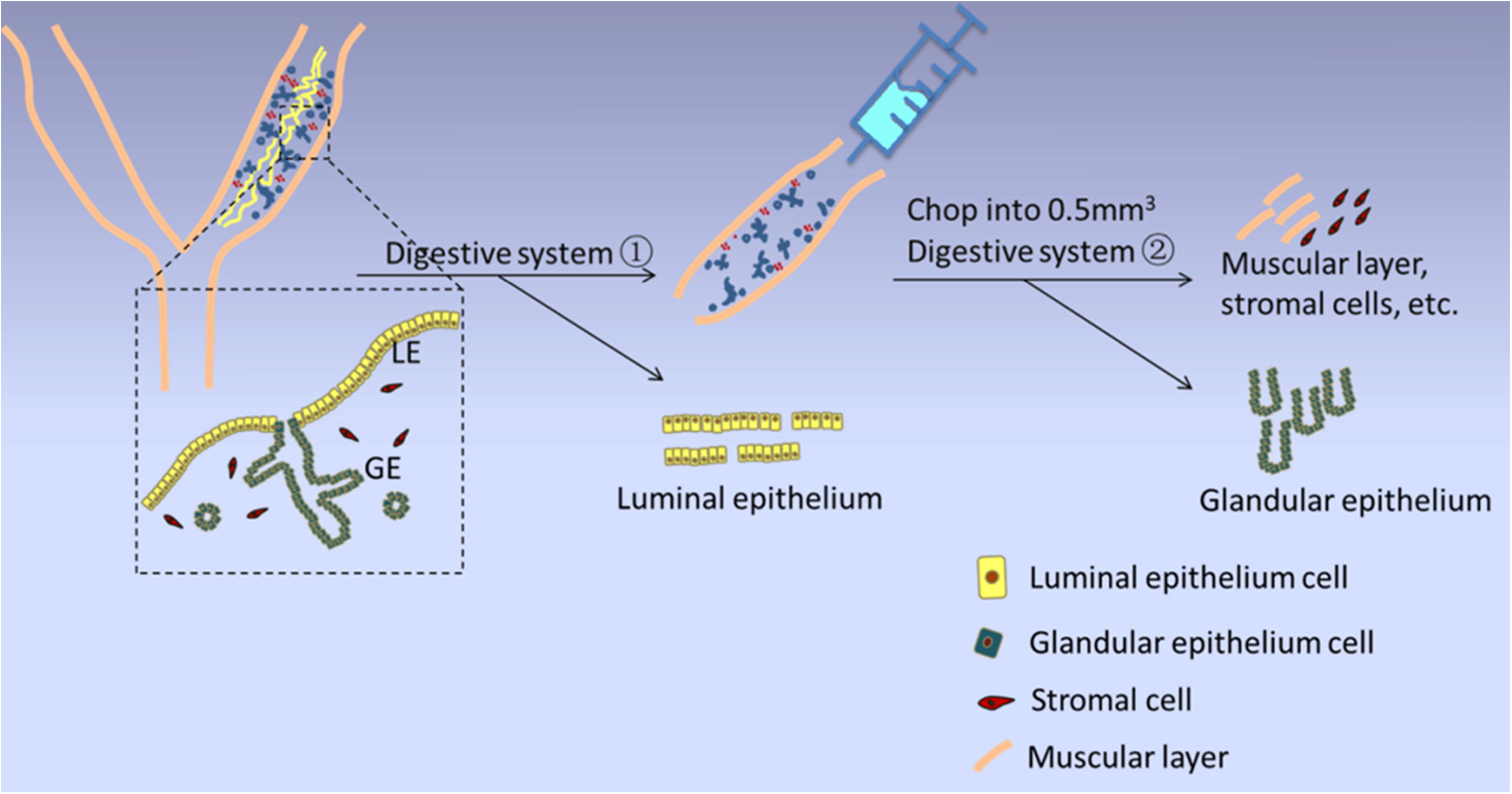
A schematic diagram depict the steps of the isolated method.

### 2. Isolation of Intact Uterine Luminal Epithelium

The critical first step in this study was the removal of the intact luminal epithelium from the uterus. We have designed the following steps to achieve this goal:

a. Digesting the entire uterine horn with appropriate trypsin concentration to lose the connection between the LE and underlying stromal tissue. Due to its complicated structure and poor digestion, the glands were still tightly attached to stromal tissue.
b. Appling a mechanical shear forced by flushing the uterine lumen with 1x HBSS to flush out the intact LE sheet. The GE and LE break down at the connecting point.

In this study, we have tested many conditions of enzyme concentration and digestion hours. We chose the following condition as the most suitable: 0.2% of trypsin digested in the uterus for 1 hour at 4 °C followed by another 1 hour at 37 °C water bath. Under this condition, the uterine LE flushed out intactly, without the glandular tissue attached, and the joint points with glands appeared as orifices instead (Fig. 2a). The E-cadherin staining in the remaining uterus showed no LE presenting, proving the LE was flushed out intactly (Fig.2b).

**Fig 2.**
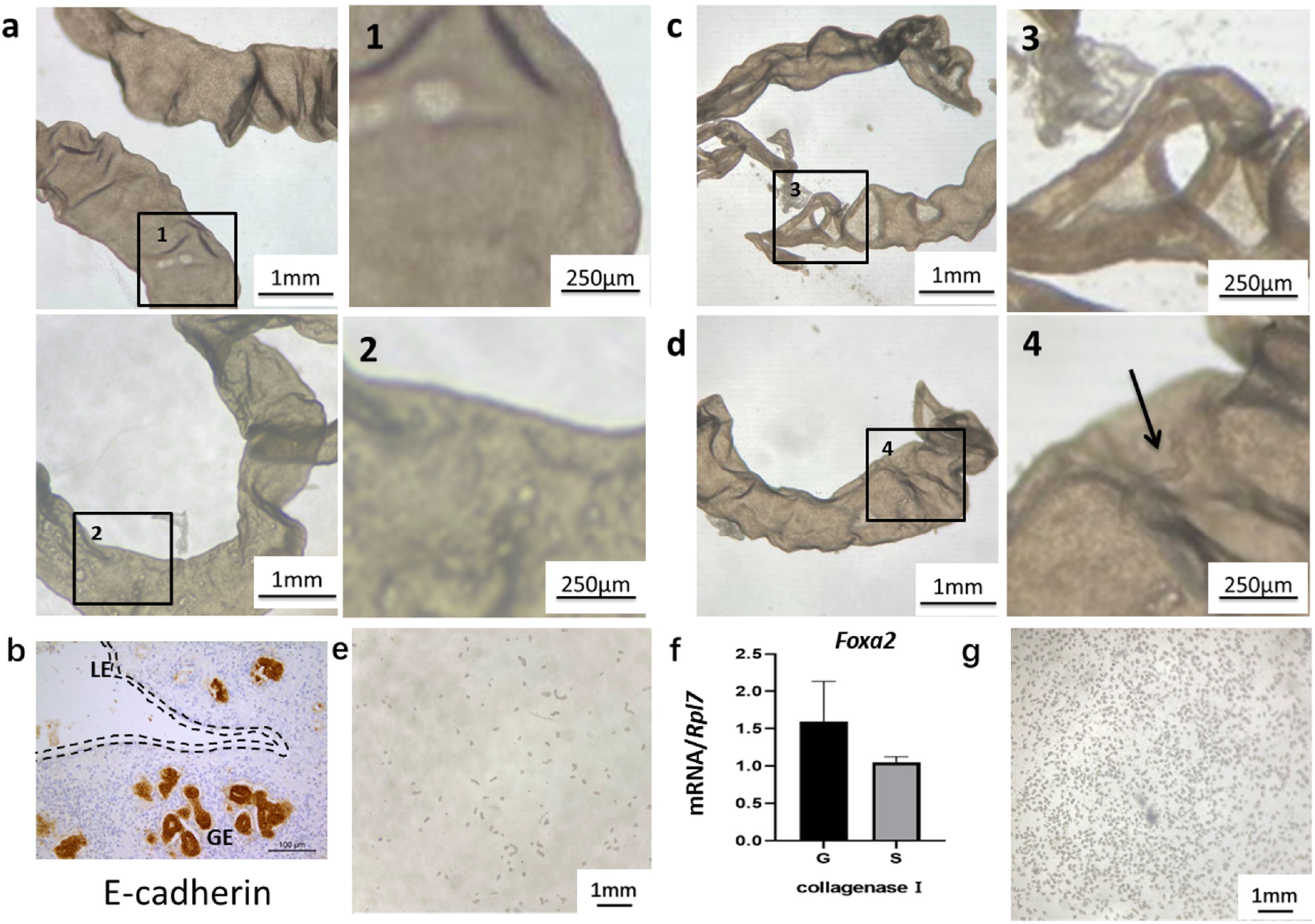
The morphology of the isolate luminal epithelium and glandular epithelium. (a) The morphology of the isolate LE. (b) E-CADHERIN staining remove LE’s uterus. The dotted line is the position of the luminal epithelium. GE: glandular epithelium; LE: luminal epithelium. (c) The morphology of the isolate LE in low trypsin concentration. (d) The isolate LE morphology in high trypsin concentration. (e) The isolate GE by collagenase I. (f) Compared the Foxa2 expression of GE and Stroma cells by collagenase I digestion. (g) The isolate GE with collagenase IV.

In case of a lower concentration of trypsin (0.1%), or shortened incubation time, in other words, incomplete digestion, stromal cells were found attached to the LE, and LE abscission is incomplete (Fig.2c). In contrast, when too much trypsin (0.3%) was used or incubated for too long, part of the glands was flushed out with the LE (Fig.2d). Both resulted in contamination of the LE fraction.

### 3. Separation of glandular epithelium out of the stroma

After removing the intact uterine LE, there were only stromal cells next to the GE cells. We then digested the remaining tissue using collagenase to free glands from the stroma. We first used Collagenase type I and Dispase II solution. However, only a few short gland fragments were recovered using these enzymes (Fig.2e). We then analyzed the gene expressions by qPCR and found that the cells from filtrate expressed a comparable level of *Foxa2*, a well-known GE marker gene, to the collected GE cells, indicating many GE cells passed through the filter and mixed with the stromal cells because of their small size (Fig. 2f). It is known that type I collagenase contains considerable tryptic activity. We hypothesized the GE might be over-digested by the tryptic activity into small pieces, even single cells, to pass through the mesh filter. Therefore, we replaced type I collagenase with type IV, which contains the lowest tryptic activity among all four types of collagenases, and repeated the separation process. The results showed that collagenase type IV digestion resulted in a good number of rounder/ellipse shape glands than that digested with type I collagenase (Fig.2g).

### 4. Purity analyses of luminal epithelium and glandular epithelium

To detect the purity of the isolated LE and GE, we tested the expression of cell type-specific markers of different epithelial cells. Calbindin-D28k is uniformly expressed in the luminal epithelial cells in the uterus of the immature mouse [23]. Herein, the mRNA expression of *Calb1* was significantly lower in the GE fraction than LE fraction but still higher when compared to the stroma fraction (Fig.3a). We further detected the immunostaining of CALB1 protein. The result showed that the whole LE, from both uterine sections and isolated fractions, highly expressed CALB1 (Fig.3g). In addition, some of the glands close to LE were also positive in CALB1 at E3.5, suggesting that CALB1 is not an excellent LE-specific marker, at least at E3.5 (Fig.3g). Furthermore, TROP2 (encoded by *Tacstd2*) has been reported to express only in uterine LE but not GE and S in adult mice [23]. We, therefore, detected the expression of *Tacstd2* mRNA by qPCR. The result showed *Tacstd2* was significantly higher expressed in the LE fraction than the GE and S (Fig.3b). This data, together with the absence of E-cadherin-positive cells on the surface of the uterine lumen after LE isolation (Fig. 2d), suggested that the intact LE was isolated, and the GE and S fractions did not contain any LE cells.

**Fig 3.**
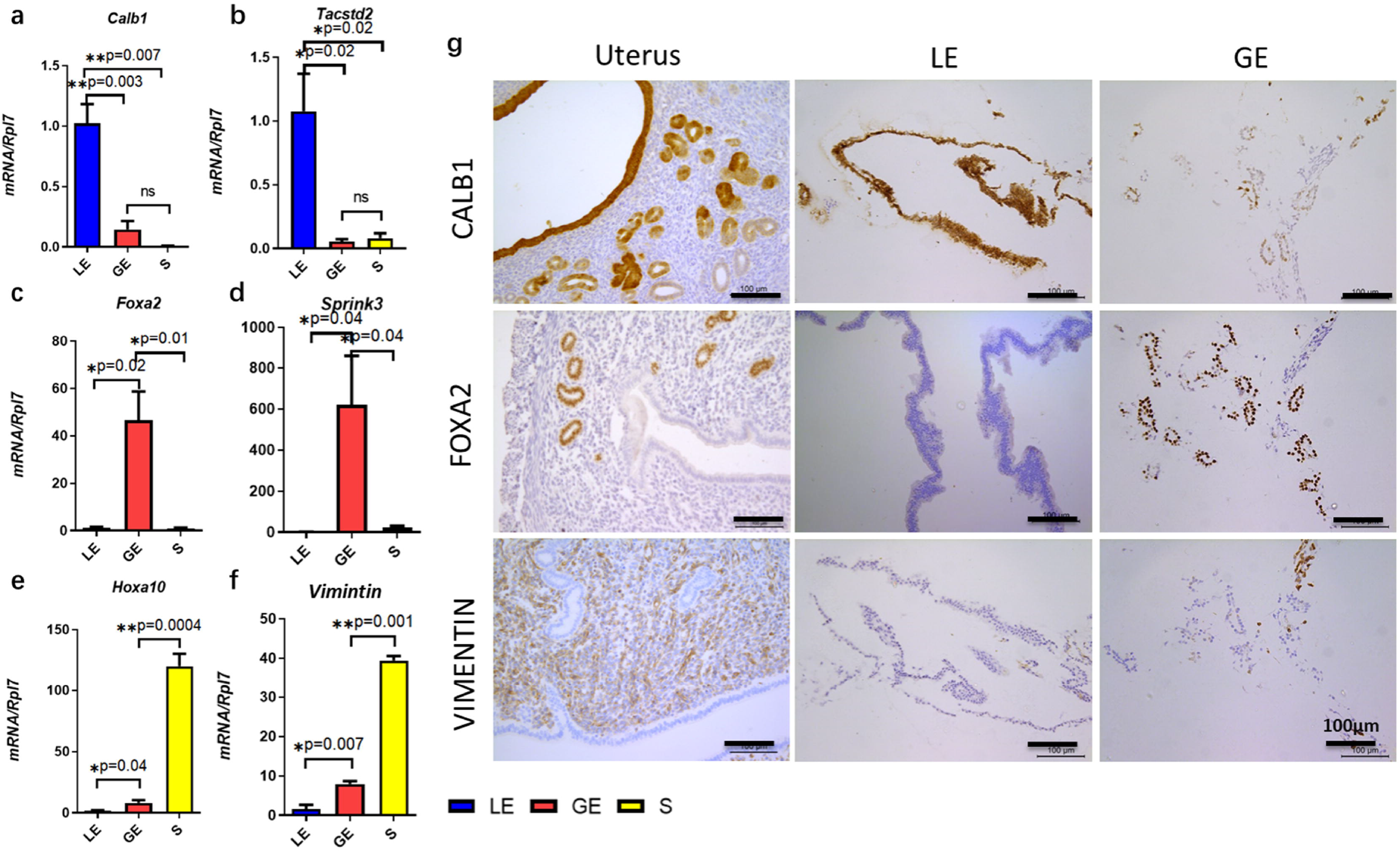
Purity analysis of LE and GE. (a, b) Quantitative PCR expression of *Calb1* and *Tacstd2*. (c, d) The mRNA levels of *Foxa2* and *Spink3* are the GE markers. (e, f) Expression of the *Hoxa10* and *Vimentin* mRNA on LE, GE, and S. (g) Representative immunofluorescence images staining for specific markers of the LE, GE, and S. Bar= 100 μm; *P < 0.05; **P < 0.01.

Next, we testted the expression of GE markers in our isolation fractions. The mRNA levels confirmed that both Foxhead box a2 (*Foxa2)* [25] and serine protease inhibitor Kazal type 3 (*Spink3*) [26] were significantly higher expressed in GE fractions compared to LE and S (Fig.3c, d). Besides, FOXA2 staining was positive in the glands of the uterine section and the isolated GE fraction but negative in the isolated LE fraction (Fig.3g). The expression of GE markers indicated that most GE cells were collected in the isolated GE fractions. We then detected the expressions of stromal cell markers Homeobox A10 (*Hoxa10*) and *Vimentin* [27, 28]. The expression of *Hoxa10* and *Vimentin* mRNA were significantly higher in the S fraction compared to the GE and LE fractions, and the immunostaining of VIMENTIN protein in the LE fraction was negative in isolated LE (Fig.3e-g). However, even, the expressions of *Hoxa10* and *Vimentin* in S were approximately 100 times and 40 times higher than in the glandular epithelium, respectively, the GE fraction still expressed a certain amount of *Hoxa10* and *Vimentin* mRNA, which were significantly higher than LE, suggesting the presence of stromal cells in the GE fraction (Fig.3e&f). The immunostaining confirmed that approximately 15% of cells were VIMENTIN positive in the GE fraction (Fig.3g), indicating the purity of the GE we isolated was around 85%. These results indicated that we achieved a high purity of isolated LE and GE fractions.

### 5. Transcriptome analysis of isolation methods

Previously studies have investigated the transcriptomic difference between the GE and LE cells on a microarray by separating the luminal and glandular epithelium from the uterus at E3.5 using LCM [22]. We, therefore, performed an RNA-Seq transcriptome analysis of the GE and LE fractions isolated in this study. Compared to the GE, there were 27756 differently expressed genes (DEGs) in LE, 5427 significantly up-regulated and 7876 significantly down-regulated, with a threshold as fold change (FC) ≥2 and p<0.05 (Fig.4a). On the other hand, we re-analyzed the raw data of GSE48239 and got 22843 DEGs, 918 significantly up-regulated and 830 significantly down-regulated, with the same threshold set up (Fig.4b). A Venn diagram showed that 1070 DEGs were overlapped between our enzyme-based RNA-Seq data (Enzyme) vs. the LCM Microarray data (LCM) (Fig.4c). A correlation analysis based on fold changes of these 1070 genes showed a significant correlation between the Enzyme and LCM data (Fig.4c). However, 11721 and 491 DEGs were still unique for either Enzyme or LCM data, respectively (Fig.4c). More than 2/3 of DEGs from the LCM data were identified in our study, while ten times more DEGs were identified uniquely in our study, which might be caused by the difference in platforms used in the two studies, the RNA-Seq used in this study is more powerful than the Microarray platform used in the LCM study. Additionally, many of the genes that were highly expressed in uterine glands (*Cxcl15, Foxa2, Prss28, Prss29, Lif, Spink3, Ttr*) and the luminal epithelium (*Wnt7a, Lpar3, Lifr, Calb1*) had been identified in our RNA-Seq data (Fig.4a). However, the volcano map showed that some of these genes did not differ significantly in the LCM-MA data (Fig.4b). For example, *Lpar3*, which encodes LPA3, the third G protein-coupled receptor for lysophosphatidic acid (LPA), is shown prominently expressed in the luminal epithelium at E3.5 and contributes to the spacing of the embryos in the uterine lumen [29]. Nevertheless, *Lpar3* was only identified as a DEG in our RNA-Seq data, not in the LCM-MA data (Fig.4a, b). These data suggested that the LE and GE isolation method we used in this study is at least comparable to the LCM method.

**Fig 4.**
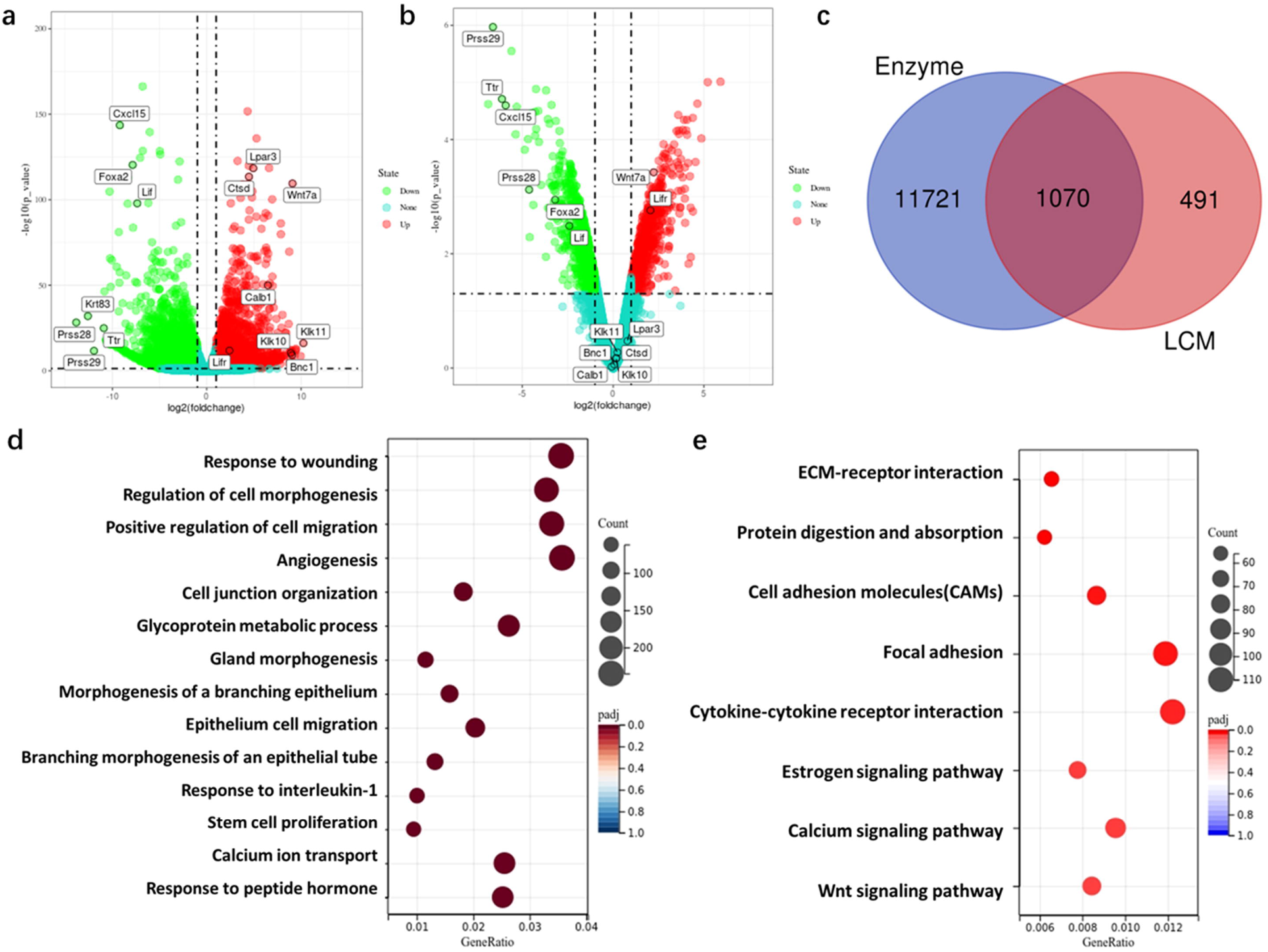
Comparing the enzyme-based method and the LCM method in transcriptome analysis. (a) Volcano plot depicting RNAseq expression analysis of the isolate LE and GE by the enzyme-based method. (b) Volcano plot showing RNAseq expression analysis of the isolate LE and GE by the LCM method. (c) Venn diagram of the different genes between LE and GE in the enzyme-based and LCM methods. (d) Gene ontology (GO) analysis relating to biological processes of the genes up-regulated in a. (e) KEGG analysis relating to the enzyme-based isolate LE and GE.

Moreover, visualization of biological process GO terms in our data showed that the DEGs were associated with gland morphogenesis, morphogenesis of a branching epithelium, epithelial cell migration, and so on (Fig.4d). KEGG analysis showed the GE and LE were different in the Estrogen signaling pathway, Wnt signaling pathway, and so on (Fig.4e). These results were associated with differences between the two epithelium types, indicating the credibility of our isolation method.

### 6. *In vitro* Culture of isolated LE and GE cells

At last, we tested if the LE and GE cells isolated in this study could be in vitro cultured for further study. We cultured LE and GE cells in both 2D and 3D conditions. In 2D culture, both luminal and glandular epithelial cells could adhere and grow on the surface of the peri dish, proving that the isolated luminal and glandular epithelial cells were viable. Besides, we tested the CALB1 and VIMENTIN expression in 2D culture by immunofluorescence. There was no cell shape VIMENTIN signal in the cultured LE, but a small number of positive cells in the cultured GE cells were consistent with our purity test results (Figs.3f&g, 5a). On the other hand, almost all the cultured LE cells expressed CALB1, only a few cultured GE cells were CALB1 positive, and all S cells were negative (Fig.5a).

**Fig 5.**
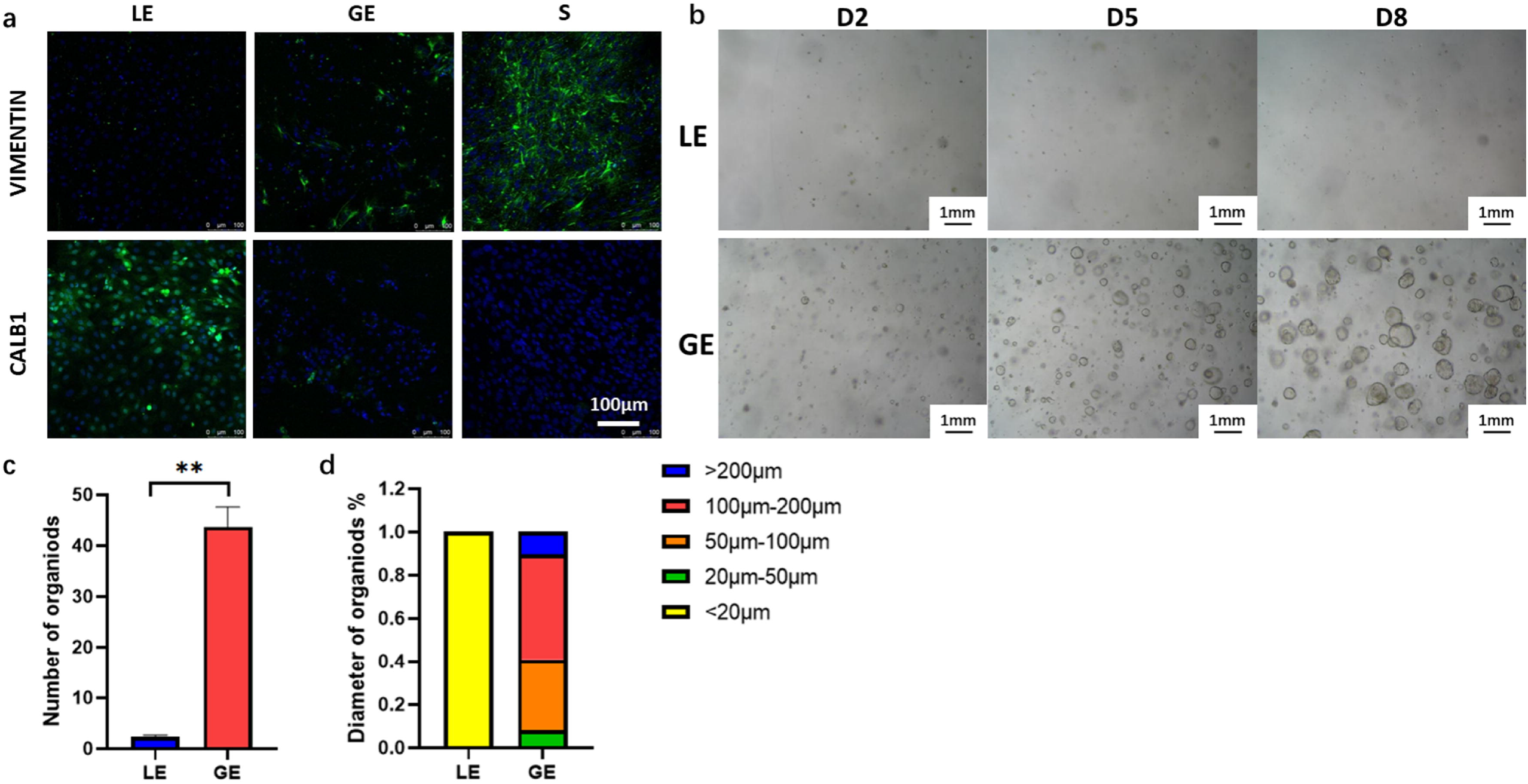
Functional verification of LE and GE cells. (a) The fluorescence intensity of CALB1 and VIMENTIN of LE and GE cells in 2D culture. (b) Recording of the LE and GE organoid culture in 8 days. The scale bar indicates 1mm. The data points represent the number (c) and diameter (d) of organoids.

Organoids are self-forming 3D reconstructions of an endometrial epithelium [30, 31]. In this study, we also cultured the organoid using our isolated luminal and glandular epithelial cells to detect their capability in 3D culture. From days 2 to 8, the GE-driven organoids gradually self-organized and grew (Fig.5b). However, only a few LE cells could form organoids (Fig.5b). The diameter and forming efficiency of organoids derived from GE were much higher than that from LE (Fig.5c, d). The diameter of LE-driven organoids was lower than 20 μm, while the diameter of the GE-organoids was most concentrated in 100-200 μm (Fig.5d). Overall, both types of isolated epithelium can maintain their viability and be cultured.

## Discussion

The endometrium comprises two distinct types of epitheliums in morphology and function: the luminal and glandular epithelium. The luminal epithelium is the first barrier of the endometrium and the place where the endometrium first contacts active embryonic trophoblast cells, which plays a vital role in embryo implantation [32]. In contrast, uterine glands provide tissue nutrition and key cytokines such as LIF for embryo implantation and development [32]. Therefore, separating these two types of epithelia is necessary to explore their different functions during pregnancy and the differentiation pattern between LE and GE during development. To our best knowledge, previously, no such technology could separate the mouse uterine LE, GE, and S cells and maintain their viability for in vitro culture. In this study, we reported a novel separation method that can simultaneously isolate the LE, GE, and S from a mouse uterus and keep the possibility of cell culture for further study.

As a cell-isolating method, purity is the most important characteristic that needs to be discussed. According to the qPCR and immunostaining results, the purity of LE isolated by our method was pure; almost no GE and S cells were found in the LE fraction. The GE fraction also contained a low level of LE, and even the immunostaining showed that the expression of CALB1 was positive in glands near the LE. The relative lower expression of *Calb1* mRNA in the GE fraction suggested that the majority of GE cells we isolated were the ones that were close to the myometrium but not near the LE. According to the immunostaining data, the GE fraction we isolated contains about 15% of S cells. The purity was not very ideal from this point of view. An additional washing step before collecting the GE fraction may help to reduce the S cell contamination but reduce the yield simultaneously. Also, the epithelial fraction isolated by other methods still contains approximately 7% to 10% of VIMENTIN-positive stromal cells [17, 33].

Previously, LCM was the only method to isolate the LE and GE from the endometrium. Filant J et al. dissected high purity of GE and LE fractions from the mouse uterus at E3.5 using LCM and then compared the transcriptomes of GE and LE cells using microarray [22]. By comparing the data of the LCM method and our enzyme-based method, we confirmed that the transcriptional profile of these two methods is similar, suggesting the purity of fractions we isolated is good enough for further study. In addition, our data showed much more DEGs than the LCM data, which can be due to the different platforms used in the two studies (RNA-Seq vs. Microarray) [22]. GO analysis refers to the DEGs between LE and GE to gland morphogenesis and morphogenesis of a branching epithelium, which were the characteristic features of glands [4]. The KEGG analysis showed that estrogen signaling was one of the most different activated pathways between the LE and GE. This pathway could be caused by the different expression of ESR1 in LE and GE cells at E3.5. Furthermore, compared to the LCM, our method was easier to provide sufficient tissue for omic analysis. Another approach to achieving separation of epithelial fraction from endometrium is pulling down the cells by immunomagnetic beads coupled with antibodies to specific cell surface proteins, e.g., EPCAM, from cell solution generated by collagenase digestion [34]. However, the lack of known specific surface proteins that can distinguish LE and GE limits the application of magnetic beads in separating these two epithelial cell types.

To verify that the epithelial cells we isolated were viable, we cultured the epithelial cells in 2D and 3D. 3D cultured organoids have been developed from various organs as powerful tools for studying tissue biology and disease. In human organoids cultured, they isolated cells by human uterine glandular epithelium. These organoids expand long-term, are genetically stable, and differentiate following treatment with reproductive hormones [31]. In mice, researchers used a mixture of luminal epithelium and glandular epithelium. Most of the organoids express glandular epithelial properties [36]. Organoids are self-organizing, genetically stable 3D culture systems containing progenitor/stem and differentiated cells resembling the origin tissue [37]. In our results, organoid cultured by glandular epithelial cells was better than luminal epithelial cells. Articles have shown that Lgr5 is broadly expressed in the uterine epithelium during embryogenesis but becomes restricted mainly to the tips of developing glands after birth, Lgr5^+^ stem/progenitor cells contributing to the development of the female reproductive tract epithelia [38]. Besides, Shafiq M. Syed suggested that Axin2 marks endometrial stem cells residing in gland bases that fuel endometrial epithelial growth and regeneration [39]. Our result also showed that there were more stem/progenitor cells in the glandular epithelium.

The endometrial epithelium experiences cyclical regression and regeneration under the regulation of ovarian steroid hormones [40]. Therefore, the 3D structure of LE changes rapidly during the estrus cycle and pregnancy days, making separating the intact LE fractions difficult. Herein, we first successfully isolated LE and GE at E3.5 because the LE is flatter in this stage compared to other stages. We also tried isolating these two epithelium types at E1.5 and E2.5. The intact LE could be removed from the uterus at these two stages (Fig.S1). However, a large number of folders due to the estrogen-caused proliferation make the LE structure more complex in the estrus stage and at E0.5 [40], and the LE isolated from these stages is often fragmented. Therefore, an additional step that manually removed the residual LE using forceps was needed before the digestion of GE out of S to avoid contamination of GE and S by LE fraction.

In summary, we provided a novel enzyme-based method to separate the endometrial LE, GE, and S cells in high amounts and purity. The cells isolated using this method could be cultured in 2D and 3D conditions, providing technical support for further cellular and molecular studies regarding the distinguishing characteristics of endometrial LE and GE. Furthermore, using our method, we compared the transcriptional profiles of LE and GE cells at E3.5, providing improved data for the gene expression difference in these two types of cells.

## Materials and methods

### Animals

All mice were housed in the SPF Experiment Animal Facility with the approval of the Institutional Animal Care and Use Committee of South China Agricultural University. Mice were maintained in a temperature- and light-controlled environment (12 h light and 12 h dark cycle) with free access to regular food and water. Mature female C57BL/6 mice (6–8 weeks old) were mated with fertile males of the same strain, and the day of the vaginal plug was marked as E0.5.

### Luminal epithelium isolation

The mouse uterus was collected after euthanasia, and the blastocyst was flushed out to confirm pregnancy at E3.5. The uterine horns have been removed the mesometrium and washed with HBSS (Sigma, SLCB9243) three times (5 mins per time). Put the uterine horns into a digestion solution containing 0.2% Trypsin and 0.6% Dispase II (Roche BR, 4942078001) in HBSS, digested at 4□ for 1 hour, and digested in a water bath at 37□ for 1 hour. After the digestion is complete, use a 10 mL syringe to draw HBSS buffer to flush out the uterine luminal epithelium.

### Glandular epithelium isolation

The remaining tissues were chopped into small cubes and enzymatically digested in 10% FBS, 0.1% Collagenase (Gibco, 17100017, 17104019), 0.6% Dispase II digestion system in a 37□, 810 rpm/30 min metal bath, add then add 1% DNase (Yeasen, 10607ES15) for 5 more minutes. After the digestion, the supernatant was passed through one 200-mesh filter, and the sieve was washed several times with an HBSS medium. The flow-through was collected and passed through 1000-mesh twice. The final filtrate contains stromal cells; the filter residue is the glandular epithelium. The sieves were inverted over a 3.5cm petri dish, and retained glandular epithelium elements were backwashed from the sieve membranes, pelleted by 300 rcf/5 min centrifugation, and collected.

### Organoid cultured

Luminal and glandular epithelial cells were centrifugated and resuspended in 75% Matrigel (CORNING, 356231)/25% DMEM/F12 (Gibco, 11039-021). 45 μL drops of Matrigel-cell suspension were plated into a 24-well plate, allowed to set at 37□, and overlaid with 750 μL organoid culture Medium [36]. The medium was changed every 2-3 d.

### Total RNA Isolation and Real-Time PCR

Total RNA was isolated using TRIZOL reagent (TaKaRa) and reverse transcripted into cDNA using HiSuperscript II kit (Vazyme, R233-01) according to the manufacturer’s introduction. Real-time PCR reactions were performed using SYBR (Vazyme, 7E381B9) method on a Bio-Rad Thermal Cycler. Expression levels were calculated by applying the comparative Cycle threshold (Ct) method.

### Immunohistochemistry (IHC)

For immunofluorescence, dissected uteri were fixed in 4% paraformaldehyde/PBS overnight at 4□. Formalin-fixed paraffin-embedded sections were deparaffinized at 55□ for an hour. Antigen retrieval was performed in a microwave in Sodium citrate solution, pH 6.0. After antigen repair, blocked for 1 h in 10% serum /PBS solution at 37□, then incubated with primary antibodies in the blocking solution overnight at 4□. The next day, sections were washed three times with PBS (5 min per wash), incubated with secondary antibodies for 1 h at 37□, and washed and incubated with HRP at 37□ for 30 min. Tissue slides were stained with diaminobenzidine (DAB) (Thermo Scientific) and counterstained with hematoxylin.

### Immunofluorescence (IF)

Cell climbing sheet fixed in 4% Paraformaldehyde for 30 min, and then permeabilized in 1% Triton X-100 (BBI Life Science, A600198)/PBS for 15 min at room temperature, blocked for 1 h in 10% Horse-serum/PBS solution, and incubated with primary antibodies in the blocking solution overnight at 4□, incubated with secondary antibodies for 30 min at 37□ and counterstained by 4, 6-diamidino-2-phenylindole dihydrochloride (DAPI) for 1 min. Cell climbing sheet imaged with confocal fluorescence scope (Leica, TCS SP8, Germany).

### RNA-Seq and Data Analysis

Tissue total RNA was extracted by The Trizol RNA Reagent (Takara, Dalian, China). The concentration and integrity of the RNA were measured by the ND-1000 Nanodrop and the Agilent 2100 TapeStation (Novogene Bioinformatic Technology, Beijing, China), respectively. The quality control parameters used in this study were: A260/A280 ratio ≥1.8, A260/A230 ratio ≥2.0, and RNA integrity number ≥8.0. cDNA libraries were generated by The TruSeq RNA sample preparation kit (Illumina, San Diego, CA, USA). RNA sequencing was performed on an Illumina HiSeq 2500 system. Raw data were processed with an in-house computational pipeline. GO, and KEGG analysis was performed using the DAVID online tools, and the false discovery rate (FDR) cutoff was set at 0.05. The RNA-seq raw data were deposited in Gene Expression Omnibus (GEO) with the accession number GSE206491.

### Statistical Analyses

Data of mice tissue were analyzed using the Student’s t-test for two-group data or one-way ANOVA with Tukey’s multiple comparisons to determine significance in three-group experiments. All statistical analyses were performed by GraphPad Prism 8 and all data were presented as Mean±SEM. Values were considered significant if P was<0.05.

## Supporting information

DEG--enzyme-based separation method

## Figure Legends

Fig S1. (a) E-CADHERIN staining of remove the LE’s uterus at E1.5 and E2.5.

