## Supplementary figures and images for "Distinguish characters of luminal and glandular epithelium from mouse uterus using a novel enzyme-based separation method"

### Supplemental Data 1

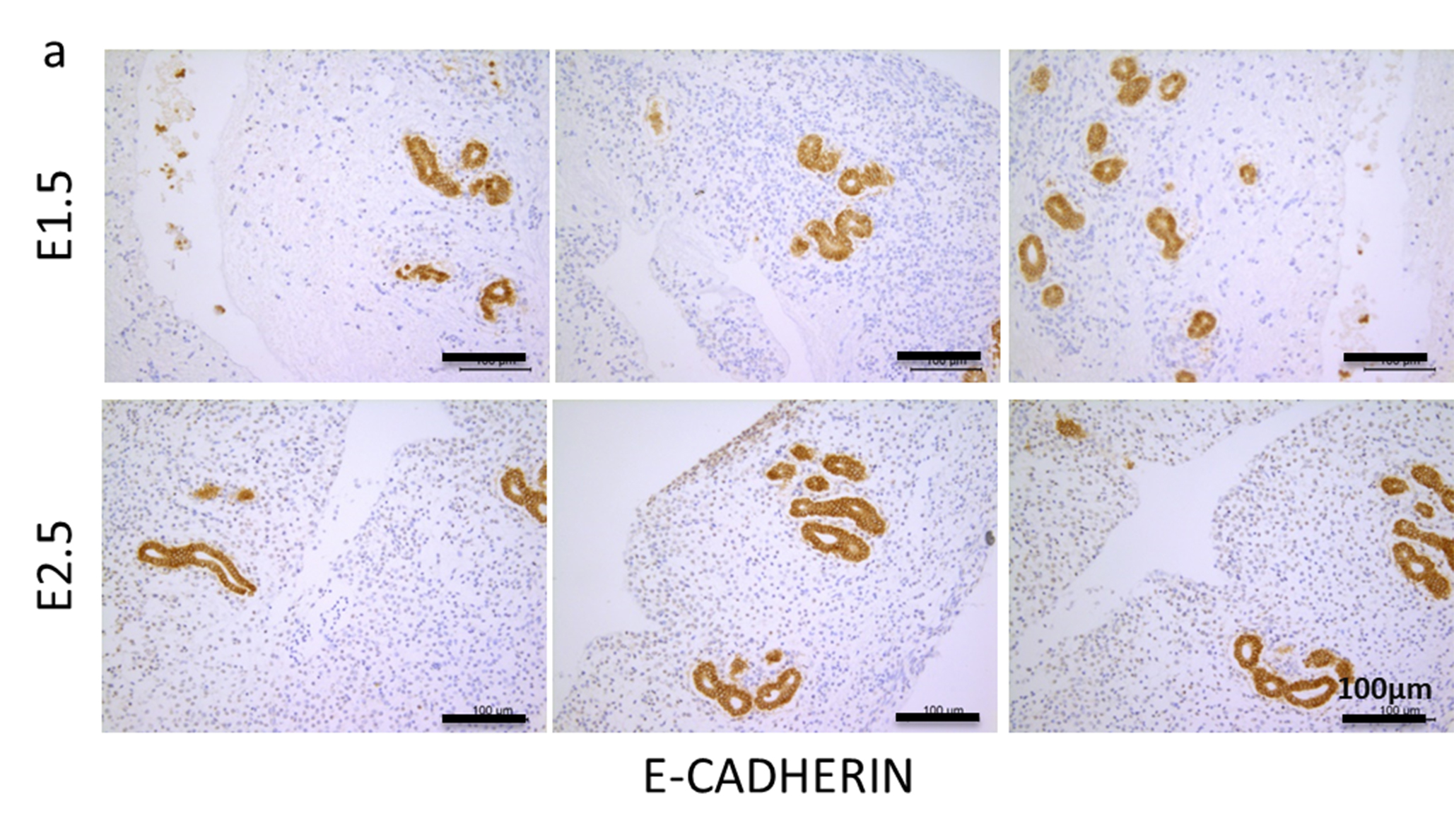
